# Early identity recognition of familiar faces is not dependent on holistic processing

**DOI:** 10.1101/287466

**Authors:** Sarah Mohr, Anxu Wang, Andrew D. Engell

## Abstract

**Highlights:**

- Holistic processing is not necessary for early identity recognition of familiar faces.
- Inverted faces and isolated features can effectively activate the perceptual memory of a familiar face.
- This effectiveness was observed for eyes, but not mouths.

**Abstract:** It is widely accepted that holistic processing is critical for early face recognition, but recent work has suggested a larger role for feature-based processing. The earliest step in familiar face recognition is thought to be matching a perceptual representation of a familiar face to a stored representation of that face, which is thought to be indexed by the N250r event-related potential (ERP). In the current face priming studies, we investigated whether this perceptual representation can be effectively activated by feature-based processing. In the first experiment, prime images were familiar whole faces, isolated eyes, or isolated mouths. Whole faces and isolated eyes, but not isolated mouths, effectively modulated the N250r. In the second experiment, prime images were familiar whole faces presented either upright or inverted. Inverted face primes were no less effective than upright face primes in modulating the N250r. Together, the results of these studies indicate that activation of the earliest face recognition processes is not dependent on holistic processing of a typically configured face. Rather, feature-based processing can effectively activate the perceptual memory of a familiar face. However, not all features are effective primes as we found eyes, but not mouths, were effective in activating early face recognition.

## 1. Introduction

Face recognition is a key facilitator of successful social interactions, as it allows for retrieval of biographical and semantic information necessary to guide appropriate social behavior (Bruce & Young, 1986; Haxby, Hoffman, & Gobbini, 2000). The earliest step in familiar face recognition is thought to be the matching of a perceptual representation of a familiar face to a stored representation of that face (Bruce & Young, 1986). This process is likely indexed by the N250r event-related potential (ERP), which is evoked by the sequentially paired presentation of a prime face (S1) followed by a target face (S2) of the same identity (Begleiter, Porjesz, & Wang, 1995; Bindemann, Burton, Leuthold, & Schweinberger, 2008; Schweinberger & Burton, 2003; Schweinberger, Pfütze, & Sommer, 1995). The N250r appears as a negative deflection over the right occipitotemporal scalp at ~200-300 ms after the onset of S2.

Although the N250r is evoked by the repetition of any pair of images of the same face, there are important differences between the response to familiar and unfamiliar individuals. The amplitude of the N250r is larger (i.e., more negative, indicating a larger repetition effect) to the repeated presentation of a familiar face than to the repeated presentation of an unfamiliar face (Begleiter et al., 1995; Dörr, Herzmann, & Sommer, 2011; Gosling & Eimer, 2011; Herzmann, Schweinberger, Sommer, & Jentzsch, 2004; Schweinberger, Huddy, & Burton, 2004; Schweinberger et al., 1995; Schweinberger, Pickering, Burton, & Kaufmann, 2002; Schweinberger, Pickering, Jentzsch, Burton, & Kaufmann, 2002). Also, unlike the N250r evoked by unfamiliar faces, the N250r evoked by familiar faces is preserved even when a small number of face stimuli intervene between the prime and the target (Dörr et al., 2011; Pfütze, Sommer, & Schweinberger, 2002; Schweinberger et al., 2004; Schweinberger et al., 2002). This modulation by familiarity distinguishes the N250r from other early face-sensitive components such as the N170 (cf. Schweinberger & Neumann, 2015).

Perhaps most compelling, the N250r occurs for familiar faces even when S1 and S2 images differ considerably in camera angle, lighting, affective state, age of the individual, haircut, etc. (Bindemann et al., 2008; Fisher, Towler, & Eimer, 2016; Kaufmann, Schweinberger, & MikeBurton, 2009; Zimmermann & Eimer, 2013). This persistence of the N250r (albeit with a diminished amplitude), despite changes in perceptual, pictorial, and structural codes, suggests that the N250r is sensitive to identity, per se, and not simple image properties (Bindemann et al., 2008; Faerber, Kaufmann, & Schweinberger, 2015; Gosling & Eimer, 2011; Schweinberger et al., 2004; Wirth, Fisher, Towler, & Eimer, 2015).

A cardinal feature of face perception is that it is thought to be primarily holistic, relying on the bound representation of all face parts, rather than a collection of individual parts (for reviews, see Richler & Gauthier, 2014; Rossion 2013). Note that holistic processing and second-order configural processing (perception dependent on the relationship *between* features within a face) are not strictly synonymous (McKone & Yovel, 2009; Piepers & Robbins, 2012), but we do not distinguish between the two in the current report. One demonstration of holistic face processing comes from the “part/whole” paradigm in which participants are asked to identify face parts of previously studied faces (Tanaka & Farah, 1993). Critically, the parts are shown either in isolation or in the context of the whole face. Participants are better able to determine the identity of the individual parts when presented in the whole face. Another classic example of the dominance of holistic processing is the face-inversion effect, which refers to a more profound performance decrement for naming faces than for naming objects when images are rotated 180° (e.g., Farah, Tanaka, & Drain, 1995; McKone 2004; Tanaka & Farah, 1993; Van Belle, De Graef, Verfaillie, Rossion, & Lefèvre, 2010; Yin 1969, but see Rezlescu, Chapman, Susilo, & Caramazza, 2016). The inversion is thought to disrupt holistic perception (Farah et al., 1995; Rossion 2008, 2009; Van Belle et al., 2010; Yin 1969), though perhaps not abolish it entirely (Richler, Mack, Palmeri, & Gauthier, 2011). These behavioral effects are reflected in early ERP components associated with face perception. Face inversion causes an increased amplitude and/or increased peak latency in the P100 (Colombatto & McCarthy, 2016; Feng, Martinez, Pitts, Luo, & Hillyard, 2012; Itier & Taylor, 2002, 2004a, 2004b) and N170 (Bentin, Allison, Puce, Perez, & McCarthy, 1996; Eimer 2000; Itier & Taylor, 2004c; Rossion et al., 1999, 2000; Wiese 2013).

If perceptual matching of facial identity drives the repetition effect, what features of the face are critical for successful matching? Similar to the P100 and N170, the N250r is affected by inversion. When S1 and S2 are both inverted, the ERP magnitude is attenuated and the onset is delayed for unfamiliar faces (Itier & Taylor, 2004a; Jacques, d’Arripe, & Rossion, 2007; Schweinberger et al., 2004; Towler & Eimer, 2016). Towler and Eimer (2016) recently suggested there is a qualitative difference in how the identity of upright and inverted unfamiliar faces is processed such that upright faces are supported by both holistic and feature-based processes, whereas inverted faces are supported solely by the latter. This is consistent with the traditional interpretation of the inversion effect as reflecting the disruption of holistic processing. However, as noted earlier, the N250r is significantly modulated by the familiarity of a face. Indeed, there are several experimental manipulations (e.g., introducing backward masks or intervening face stimuli) that reduce or abolish the N250r evoked by unfamiliar faces, but have little effect on the N250r evoked by familiar faces (Dörr et al., 2011; Pfütze et al., 2002; Schweinberger et al., 2004; Schweinberger et al., 2002). Moreover, there is evidence that familiarity with a face facilitates feature-based processing (di Oleggio Castello, Wheeler, Cipolli, & Gobbini, 2017). Therefore, the dependence on holistic processing might not generalize to familiar faces.

In the current studies, we investigated whether holistic processing is necessary for early visual recognition of familiar faces, as indexed by the N250r. In the first study, S1 depicted either full faces, isolated eyes, or isolated mouths. A significant N250r to a target face preceded by a face-part prime would suggest successful activation of the visual representation of a face without benefit of a full-face context (i.e., holistic). In the second study, S1 was presented either upright or inverted. In all cases, S2 was a full upright face. This is a second critical difference (the first being the use of familiar faces) between the current work and prior studies that inverted both S1 and S2 (Itier & Taylor, 2004a; Jacques et al., 2007; Schweinberger et al., 2004; Towler & Eimer, 2016). In the current work, a repetition effect requires that an upright target face be matched to the perceptual memory evoked by the inverted prime face. Such a repetition effect would suggest that the perceptual memory of the identity was successfully extracted from the inverted prime and therefore that the earliest stages of face recognition are not dependent on holistic processing.

## 2. Methods

### 2.1. Participants

Participants were recruited from the Kenyon College campus and surrounding community and compensated for their participation. All participants had normal or corrected-to-normal vision. After completion of the EEG study, participants were asked to indicate their familiarity with each of the famous faces used in the experiment. Participants who were insufficiently familiar with the faces (unable to identify at least 75% of the individuals) were excluded from analysis. All participants gave written and informed consent. The Kenyon College Institution Research Board approved this protocol.

#### 2.1.1. Study 1: Face parts

A total of 35 individuals (20 female, 15 male) aged between 18 and 59 (*M* = 21.0) participated in Study 1. Two participants were excluded due to excessive noise in the EEG data. Nine participants were excluded from analysis due to insufficient familiarity with the famous faces. Thus, there were 24 participants (15 female, 9 male) aged between 18-22 (*M* = 20.2) in the included in the analyses.

#### 2.1.2. Study 2: Inversion

A total of 32 individuals (24 female, 8 male) aged between 18 and 23 (*M* = 20.1) participated in Study 2. Two participants were excluded due to excessive noise in the EEG data. Three participants were excluded from analysis due to insufficient familiarity with the faces.

Two participants were unable to complete the post-test familiarity evaluations due to technical issues. These participants were *not* excluded from the analysis, which potentially increases the probability of Type II, not Type I error. Inclusion of two participants with insufficient familiarity with the faces (if that were the case) would diminish the likelihood of finding a result predicated on familiarity. Thus, there were 27 participants (22 female, 5 male) aged between 18-23 (*M* = 20.2) in the included in the analyses.

### 2.2. Stimuli

Of the 1580 identities within the MSRA-CFW: Data Set of Celebrities Faces on the Web (Zhang, Zhang, Wang, & Shum, 2012), 220 were selected as being most likely to be known to college students. In an independent experiment, Kenyon College students ranked the familiarity of these 220 individuals and the seventy-two most familiar celebrities were selected for use in the current experiments. Two pictures of each celebrity were downloaded from Google Images. The image pairs were selected so that they differed in a meaningful way such as facial expression, hairstyle, age, or shooting angle. Pictures were cropped to isolate the face and resized to be 500 pixels tall with a resolution of 72 pixels per inch. Image width varied as a function of each face’s natural shape. For each image, a version was created in which the eyes or the mouth were isolated. To do so, the scene and aperture filters in Photoshop CS6 were used to blur the surrounding face (see <u>Figures 1</u> and <u>2</u>; FIRST ROW).

**Figure 1:**
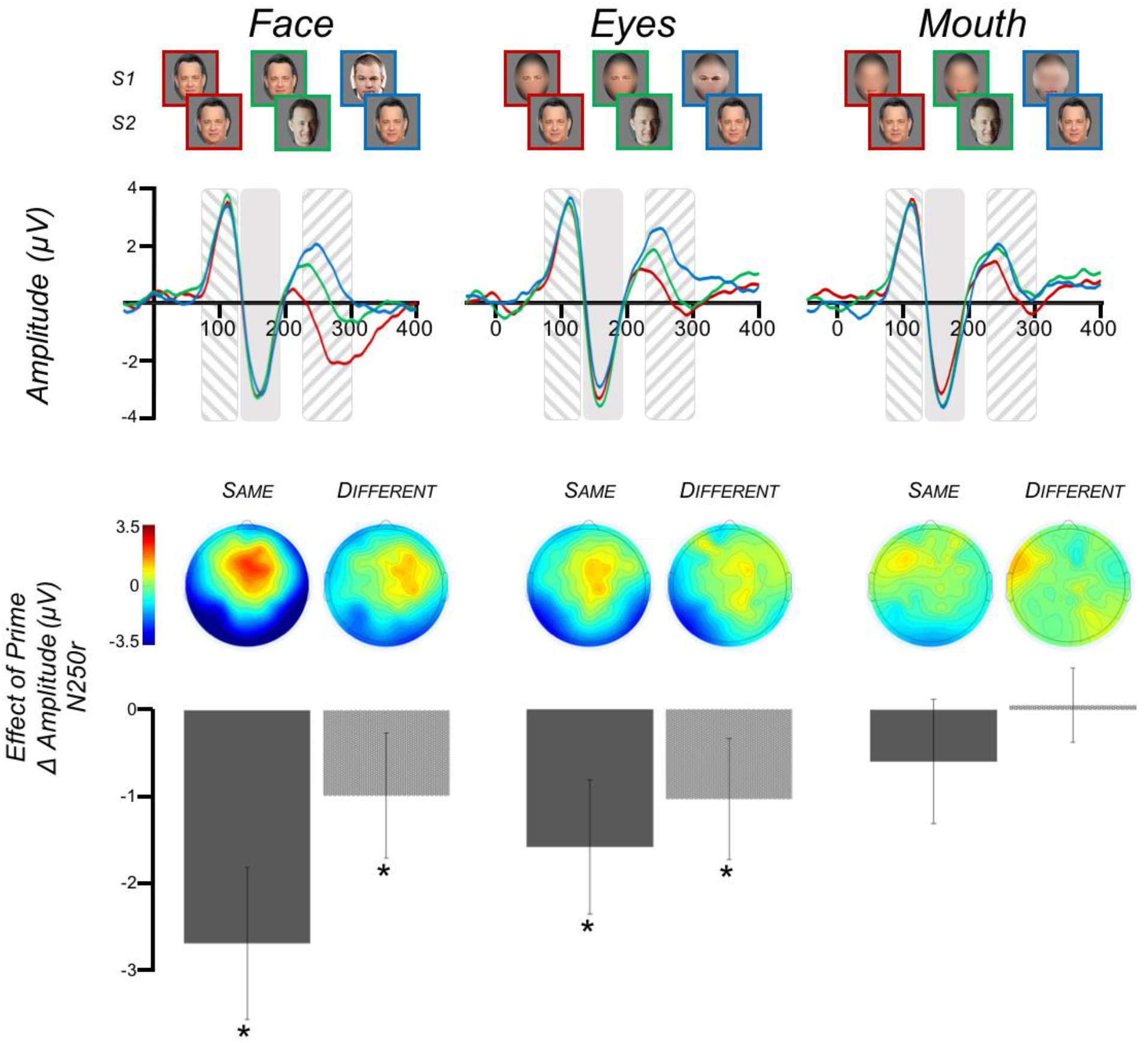
Study 1 Stimuli and ERP results. (FIRST ROW) The nine trial types used in Study 1 were: three levels of prime (same, different, none) x three levels of feature (faces, eyes, mouth). (SECOND ROW) Grand-average ERPs (N=24) across electrodes P_8_, P_10_, TP_8_, P_7_, P_9_, TP_7_. The time-period of the P100 is indicated by the area with downward slashes. NOTE: this figure displays the waveform across the electrodes listed above, whereas the P100 statistics were done on the average waveform of electrodes C_1_, C_z_, C_2_. The time-period of the N170 is indicated by the solid shaded area. The time-period of the N250r is indicated by the area with upward slashes. The waveforms are color-coded such that RED is the prime_same_ condition, GREEN is the prime_different_ condition, and BLUE is the prime_none_ condition. (THIRD ROW) Topographical maps of the N250r priming effect at each cell of the feature factor. The “Same” maps display the prime_same_ vs. prime_none_ simple contrast, whereas the “Different” maps display the prime_different_ vs. prime_none_ contrast. (FOURTH ROW) N250r mean amplitude differences across electrodes P_8_, P_10_, TP_8_, P_7_, P_9_, TP_7_. The contrasts displayed are the same as for the topographical maps. Error bars indicate the 95% confidence interval.

**Figure 2:**
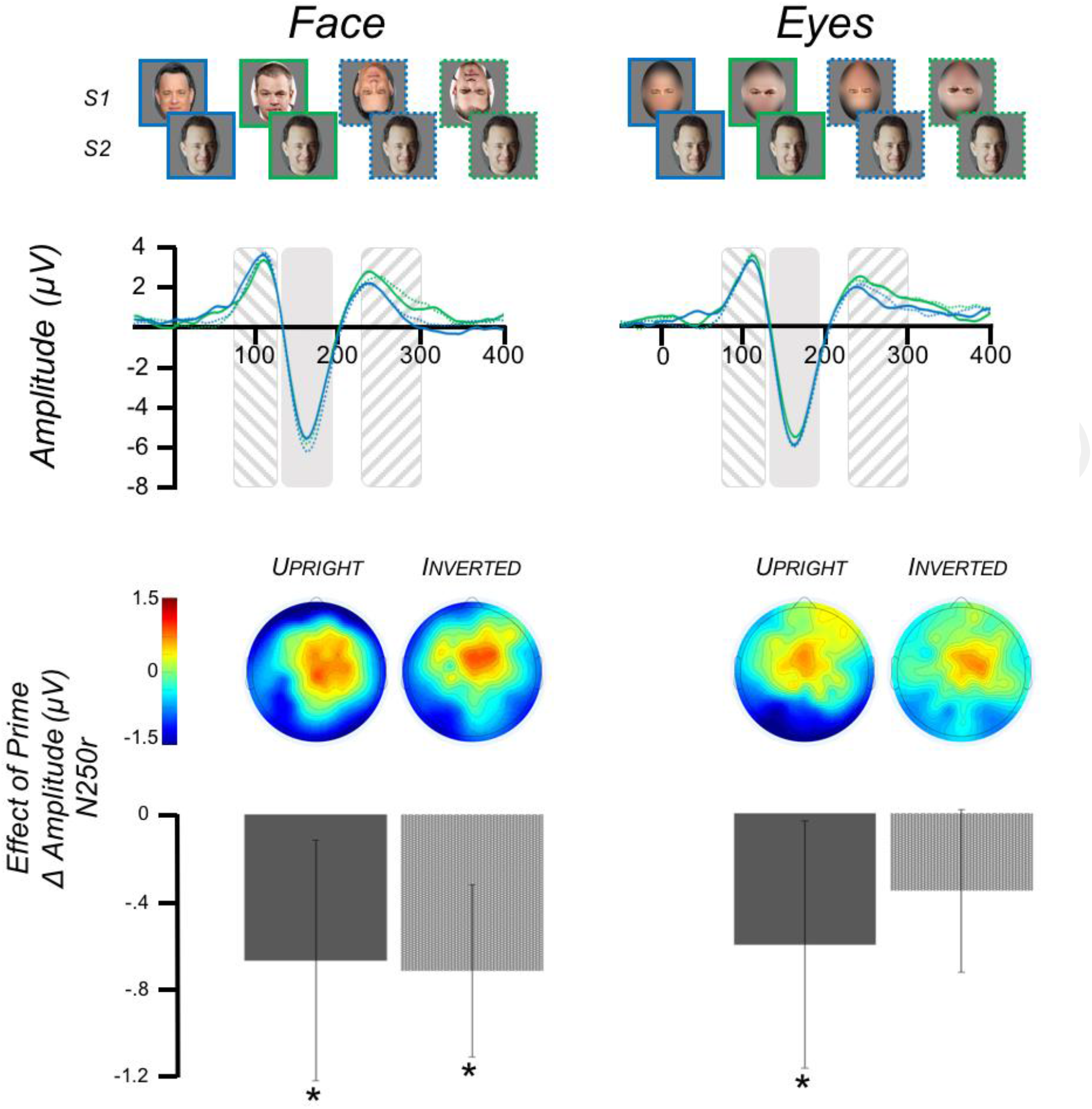
Study 2 Stimuli and ERP results. (FIRST ROW) The eight trial types used in Study 2 were: two levels of prime (none, different) x two levels of feature (face, eyes) x two levels of orientation (upright, inverted). (SECOND ROW) Grand-average ERPs (N=27) across electrodes P_8_, P_10_, TP_8_, P_7_, P_9_, TP_7_. The time-period of the P100 is indicated by the area with downward slashes. NOTE: this figure displays the waveform across the electrodes listed above, whereas the P100 statistics were done on the average waveform of electrodes C_1_, C_z_, C_2_. The time-period of the N170 is indicated by the solid shaded area. The time-period of the N250r is indicated by the area with upward slashes. The waveforms are color-coded such that SOLID BLUE is the prime_diff_orientation_up_ condition, SOLID GREEN is the prime_none_orientation_up_ condition, DASHED BLUE is the prime_diff_orientation_inv_ condition, and DASHED GREEN is the prime_none_orientation_inv_ condition. (THIRD ROW) Topographical maps of the N250r priming effect. The maps display the prime_diff_ vs. prime_none_ simple contrast for each cell of the feature and orientation factors. (FOURTH ROW) N250r mean amplitude differences across electrodes _P8_, P_10_, TP_8_, P_7_, P_9_, TP_7_. The contrasts displayed are the same as for the topographical maps. Error bars indicate the 95% confidence interval. Asterisks indicate differences significantly priming effects, *p* < .05.

### 2.3. Experimental Procedure

Stimulus presentation was controlled by PsychoPy (Peirce 2007) and displayed on a 27” LCD display with a resolution of 1920 × 1080 and a refresh rate of 60 Hz. Participants were seated ~70 cm from the display; the exact distance varied slightly to accommodate participant comfort.

The stimulus presentation was modeled after the paradigm described in Schweinberger and colleagues (2002). All trials began with the presentation of a black fixation cross on a grey background displayed for 500 ms, followed by presentation of the prime face (S1) for 500 ms. S1 was replaced by a small green circle displayed in the center of the screen for 1300 ms, followed by presentation of the target face (S2) for 1250 ms. The location of S2 was offset from the location S1 to decrease any effects of retinal and/or low-level visual adaptation. The randomly chosen offset was either horizontal (48 pixels), vertical (27 pixels), or diagonal (55 pixels). Participants were asked to indicate the sex of the person depicted in S2 by pressing one of two buttons as quickly as possible after face onset. The following trial began after a 2450 ms interstimulus interval.

#### 2.3.1. Study 1: Face Parts

There were nine variations of the S1 and S2 pairings in this 3 × 3 design (<u>Figure 1</u>; FIRST ROW). The “prime” factor included levels of prime-same (prime_same_), prime-different (prime_different_), and unprimed control (prime_none_). In the prime_same_ condition, S1 and S2 were the same picture of the same familiar identity (e.g., Paul Newman followed by the same picture of Paul Newman). In the prime_different_ condition, S1 and S2 were different pictures of the same familiar identity (e.g., Paul Newman followed by a different picture of Paul Newman). Lastly, in the prime_none_ condition, S1 and S2 were different familiar identities (e.g., Paul Newman followed by Robert Redford). These conditions were fully crossed with the three levels of the “feature” factor: full face (feature_face_), eyes (feature_eyes_), mouth (feature_mouth_). There were 18 trials of each of the 9 conditions. These 162 trials were presented randomly across 18 counterbalanced blocks.

#### 2.3.2. Study 2: Inversion

There were eight variations of the S1 and S2 pairings 2 × 2 × 2 design (<u>Figure 2</u>; FIRST ROW). The “prime” factor included levels of prime-different (prime_different_) and unprimed control (prime_none_). So unlike Study 1, in Study 2 all prime stimuli were different images of the same identity. The “feature” factor included levels of full face (feature_face_) and eyes (feature_eyes_). An “orientation” factor added in this study included levels of upright (orientation_up_) and inverted (orientation_inv_), the latter being a face rotated 180°. There were 24 trials for all of the face conditions. Due to a coding error, there was a slight imbalance in the number of eye trials such that upright eyes (primed or unprimed) were presented 22 times and inverted eyes (primed or unprimed) were presented 26 times. The 192 trials were presented randomly across 24 counterbalanced blocks.

### 2.4. EEG Acquisition and Analysis

Continuous biopotential signals were recorded using the ActiveTwo BioSemi amplifier system (BioSemi, Amsterdam, The Netherlands). EEG was acquired from 64 scalp electrodes arranged in the 10/20 system. Two external electrodes were placed on the mastoids to be used as an offline reference. Two external electrodes were placed approximately 1 cm lateral and 1 cm inferior to the outer canthus of the left eye to record the horizontal and vertical electrooculogram (EOG), respectively.

All signals were digitized and recorded on an Apple Mac Mini running ActiView software (BioSemi) at a sampling rate of 2048 Hz. Off-line analysis was conducted with the EEGLAB (Swartz Center for Computational Neuroscience, La Jolla, CA, USA) MATLAB toolbox and the ERPLAB plugin (Steve Luck, UC-Davis Center for Mind and Brain, Davis, CA, USA).

EEG data were imported with an initial reference of the averaged mastoids, downsampled to 256 Hz, and bandpass filtered with half-amplitude cutoffs of 0.5 - 100 Hz. Epochs event-locked to the onset of S2 were extracted from the continuous EEG (−1000 to 2000). ICA decomposition was run to identity eye-movement, blink, and other artefactual components, which were removed from the data. The data were then rereferenced to the mean of all scalp electrodes.

ERPs were generated by averaging the EEG epochs from each electrode for each experimental condition. ERPs were baseline normalized by subtracting the average of a 150 ms prestimulus epoch from each time point and were lowpass filtered with a 2^nd^ order Butterworth filter with a half-amplitude cutoff of 40 Hz. The grand average ERP waveform was produced by averaging across all participants’ ERPs.

Electrodes and the time-windows for analysis were selected to investigate three components of interest: P100, N170, and N250r. These electrodes and time windows were selected based on inspection of the experiment-wide grand-average ERP and prior literature that used the same, or very similar, parameters (Herzmann 2016; see Schweinberger & Neumann, 2015 for a review; Schweinberger et al., 2004). Mean amplitudes were calculated for the P100 across electrodes C_1_, C_z_, and C_2_ from 75-125 ms. Our analysis of the N170 (130 – 190 ms) and N250r (225 – 300 ms) focused on electrodes P_8_, P_10_, TP_8_ in the right hemisphere and P_7_, P_9_, TP_8_ in the left hemisphere.

Mean amplitudes for Study 1 were analyzed using a repeated-measures ANOVA with within-subjects factors of hemisphere (hem_right_, hem_left_), prime (prime_same_, prime_different_, prime_none_) and feature (feature_face_, feature_eyes_, feature_mouth_). The Greenhouse-Geisser correction was used to correct for any violations of sphericity. Mean amplitudes for Study 2 were analyzed using a repeated-measures ANOVA with the within-subject factors of hemisphere (hem_right_, hem_left_), prime (prime_different_, prime_none_), feature (feature_face_, feature_eyes_), and orientation (orientation_up_, orientation_inv_). For both studies, the hemisphere factor was only included in the analyses of the N170 and N250r. Main and interaction effects were explicated with paired-samples *t*-tests.

## 3. Results

### 3.1. Study 1: Face Parts

#### 3.1.1. P100

There were no main or interaction effects of prime or feature (*p*s > .05).

#### 3.1.2. N170

There were no main or interaction effects of hemisphere, prime, or feature (*p*s > .05).

#### 3.1.3. N250r

There was no effect of hemisphere (*p* > 0.05). There were significant main effects of prime [*F*(1.55, 35.58) = 22.80, *p* < 0.001, 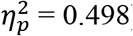] and feature [*F*(1.44, 33.13) = 17.42, *p* < 0.001, 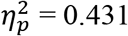]. These main effects were qualified by a significant prime × feature interaction [*F*(3.31, 76.21) = 7.25, *p* < 0.001, 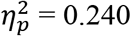] (<u>Figure 1</u>), and a significant hemisphere × prime interaction [*F*(1.69, 38.85) = 3.61, *p* = .044, 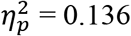]. There was no three-way interaction of hemisphere × prime × feature (*p* > .05)

To better understand the prime × hemisphere interaction, we used paired-sampled *t*-tests to evaluate the priming effect (i.e., prime_same_ vs. prime_none_, and prime_different_ vs. prime_none_) within each hemisphere. Within the left hemisphere, a significant priming effect was evoked by both prime_same_ (mean difference: −1.75 ± .33 μV, *p* < .001) and prime_different_ (mean difference: −1.03 ± .26 μV, *p*= .001), whereas only prime_same_ (mean difference: −1.51 ± .32 μV, *p* < .001) evoked an effect in the right hemisphere.

To better understand the prime × feature interaction, we used paired-sampled *t*-tests to evaluate the priming effect (i.e., prime_same_ vs. prime_none_, and prime_different_ vs. prime_none_) at each of the three feature levels (feature_face_, feature_eyes_, feature_mouth_) (see <u>Figure 1</u>, FOURTH ROW). At the feature_face_ level a significant N250r was evoked by both prime_same_ (mean difference: −2.69 ± .43 μV, *p* < .001) and prime_different_ (mean difference: −0.98 ± .35 μV, *p* = .010). At the feature_eye_ level a significant N250r was evoked by both prime_same_ (mean difference: −1.59 ± .37 μV, *p* < .001) and prime_different_ (mean difference: −1.03 ± .34 μV, *p*= .006). At the feature_mouth_ level there were no significant priming effects (*p*s > .05).

Finally, we directly contrasted the prime effect of faces with the prime effect of eyes with a paired-samples *t*-test of prime_same_ – prime_none_, and prime_diff_ - prime_none_ conditions for each of the features. There was a significantly greater effect for same faces than same eyes (mean difference: −1.10 ± 1.93 μV, *p* = .011), but no difference in effect for different faces and different eyes (mean difference: .05 ± 1.64 μV, *p* > .05).

### 3.2. Study 2: Inversion

#### 3.2.1. P100

There were no main or interaction effects of prime, feature, or inversion (*p*s > .05).

#### 3.2.2. N170

Inverted faces evoked a larger N170 than upright faces [*F*(1, 26) = 6.57, *p* = 0.017, 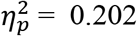]. There were no other main or interaction effects (ps > .05).

#### 3.2.3. N250r

There were significant main effects of hemisphere [*F*(1, 26) = 9.27, *p* = 0.005, 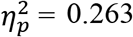] and priming [*F*(1, 26) = 22.60, *p* < 0.001, 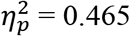]. There were no other main or interaction effects (*p*s > .05) indicating that the strength of the priming effect did not vary as a function of feature or orientation (see <u>Figure 2</u>, FOURTH ROW).

## 4. Discussion

Here we report two studies that support the notion that early face recognition is not solely dependent on holistic processing, but rather can be successfully activated by feature-based processing. However, we found that only eyes effectively engaged early recognition systems, suggesting that all features are not equipotent. These findings serve to deemphasize the role of holistic processing in familiar face perception and, in the context of the extant literature, suggest a prominent role of experience in the processes recruited to support face perception.

### 4.1. Is holistic processing necessary for facial recognition?

The results of both studies suggest that activation of a stored identity representation can occur without the benefit of holistic processing.

In Study 1, we disrupted holistic processing by presenting a single isolated facial feature (eyes or mouth) that was largely devoid of the configural information used in holistic processing (see below for a discussion of the potential effect of residual configural information). We found that isolated eyes effectively activated the perceptual memory of the prime face, as indexed by the N250r component evoked by the target face. This suggests that feature-based processing is sufficient for engaging the perceptual memory thought to be the first step in familiar face recognition. However, trials with mouth primes did not modulate the N250r regardless of whether the mouth was isolated from the same or a different picture as the target. The implication of the privileged nature of eyes is discussed in greater detail in section 4.3.

To further test the feature-based processing effect found in Study 1, we ran a second study that used a different technique for disrupting holistic processing. In Study 2, inverted faces, which are known to disrupt holistic processing (Farah et al., 1995; Rossion 2008, 2009; Van Belle et al., 2010; Yin 1969), were used as prime stimuli. If holistic processing is necessary for recognition, one would expect an interaction between prime (primed or unprimed) and orientation (upright, inverted), such that the N250r for primed vs. unprimed faces would only be observed in the upright condition. To the contrary, we found no interaction between these factors (see <u>Figure 2</u>, THIRD ROW). Moreover, pairwise comparisons found that the N250r was significant for prime trials of upright faces, upright eyes, and inverted faces, though inverted eyes did not reach significance. Put plainly, the orientation of a face had no effect on the activation of the perceptual memory of the face’s identity as indexed by the N250r.

Together, these studies suggest that holistic processing is not necessary to activate perceptual memories of facial identity of familiar faces, which is consistent with behavioral work suggesting that holistic processing is not strongly correlated with face recognition abilities (Konar, Bennett, & Sekuler, 2009; Rezlescu, Susilo, Wilmer, & Caramazza, 2017; Richler, Floyd, & Gauthier, 2015; Sunday, Richler, & Gauthier, 2017, but see DeGutis, Wilmer, Mercado, & Cohan, 2013; Wang, Li, Fang, Tian, & Liu, 2012).

It is important to note that the current results do not preclude a role for holistic processing in face recognition. Though we propose that it is unnecessary for producing the N250r, it is likely functioning in parallel. Towler and Eimer (2016) reported that while internal or external face features evoked the N250r, there was a super-additive effect for full faces. That is, the magnitude of the N250r to full face primes was larger than the sum of the effect to internal and external feature primes, which implies a unique contribution from holistic processing. Consistent with this, we found that while eyes evoked a significant repetition effect, the effect was greater for full faces. But this was only true when the prime and target were of the same image, as was the case in Towler and Eimer (2016). When the prime and target were of different images there was no difference in the strength of the face and eye priming effects. This raises the possibility that the contribution of holistic processing observed in that prior report was owed to pictorial similarity and low-level features.

It is important to point out that the manipulation in Study 1 did not entirely obscure second-order configural information because the blurred portions of the face retained some configural information (see <u>Figure 1</u>). However, it is unlikely that this meaningfully contributed to the N250r because it was observed for eyes, but not for mouths. If the preserved configural information was responsible for the N250r we would expect to see the effect in both eye and mouth conditions.

### 4.2. The role of familiarity in shaping face processing

We found that the N250r evoked by inverted primes did *not differ* from upright primes. This result is in contrast to prior work that found upright faces served as more potent primes than inverted faces (Itier & Taylor, 2004a; Jacques et al., 2007; e.g., Towler & Eimer, 2016). However, there are two important features that differentiate this study from prior work.

First, and most important, we used familiar faces as opposed to unfamiliar faces. This is a critical distinction because the effects evoked by unfamiliar faces might not generalize to familiar faces. Repetition of familiar faces results in a larger (Begleiter et al., 1995; Dörr et al., 2011; Gosling & Eimer, 2011; Herzmann et al., 2004; Schweinberger et al., 2004; Schweinberger et al., 1995; Schweinberger et al., 2002; Schweinberger et al., 2002) and more robust (Dörr et al., 2011; Pfütze et al., 2002; Schweinberger et al., 2004; Schweinberger et al., 2002) N250r than repetition of unfamiliar faces. Moreover, Burton and colleagues (2015) have recently made a strong case, based primarily on behavioral studies, against a primary role of configural processing in familiar face recognition. Though the terms configural and holistic processing are often used interchangeably, they are not synonymous (Maurer, Grand, & Mondloch, 2002; Piepers & Robbins, 2012; Richler & Gauthier, 2014). Nonetheless, both require integration of more than one face-part and can thus be similarly contrasted to feature-based processing.

Second, the current study is the first to investigate the N250r response to an upright face that has been primed by an inverted face. Prior studies presented both the prime and the target face as upright or inverted. Therefore, any modulation of the ERP evoked by the target faces in the current study can be attributed solely to the influence, if any, of the previously presented prime. That is, our design removes potential confounds introduced by presenting an inverted target face.

Considering our finding in the context of the prior literature, we support the notion that familiarity with faces is associated with a diminished importance of holistic processing during recognition (see also di Oleggio Castello et al., 2017).

### 4.3. Significance of Eyes in Face Recognition

Eyes seem to have a privileged role in feature-based processing, as we found that eyes, but not mouths, were effective primes. This privileged role is not entirely unexpected given that eyes are perhaps the most important feature in face perception. They convey critical information regarding affective state (Schyns, Petro, & Smith, 2007), attentional focus (Langton, Watt, & Bruce, 2000), and identity (Schyns, Bonnar, & Gosselin, 2002). It is therefore not surprising that deficits in typical social behavior such as Autism are often associated with impairments in eye processing (see Itier & Batty, 2009 for a review). ERP studies have found that the amplitude and latency of the face-selective N170 is modulated by eyes presented in isolation rather than in the context of a full face (Bentin et al., 1996; Itier, Latinus, & Taylor, 2006; Nemrodov & Itier, 2011; Zhang, Sun, & Zhao, 2017). Moreover, results from an intracranial EEG study of face and eye perception show that eye-sensitive regions are more abundant than face-sensitive regions throughout the ventral occipitotemporal face processing system (Engell & McCarthy, 2014).

### 4.4. Conclusions

In summary, the current study provides evidence that familiar face recognition is not solely dependent on holistic processing. Feature-based processing can effectively activate the perceptual memory of a familiar face. Compared to prior work that focused on unfamiliar faces, these results suggest an increased emphasis on facial features as a function of familiarity. However, all features are not created equally. We found that only eyes, but not mouths, were effective in activating early face recognition. This is likely owed, in large part, to the essential social information conveyed by the eyes.

## 5. Acknowledgments

We thank Henry Quillian, Claire Robertson, Josue Parr, Samantha Montoya, and Anoohya Muppirala for help in data collection, and Grit Herzmann for her comments on an earlier draft of this manuscript. This research was funded, in part, by the Kenyon Summer Science Program.

